# Genetic variance in fitness and its cross-sex covariance predict adaptation during experimental evolution

**DOI:** 10.1101/2020.02.26.966119

**Authors:** Eva L. Koch, Sonja H. Sbilordo, Frédéric Guillaume

## Abstract

In presence of rapid environmental changes, it is of particular importance to assess the adaptive potential of populations, which is mostly determined by the additive genetic variation (V_A_) in fitness. In this study we used *Tribolium castaneum* (red flour beetles) to investigate its adaptive potential in three new environmental conditions (Dry, Hot, Hot-Dry). We tested for potential constraints that might limit adaptation, including negative genetic covariance between female and male fitness. Based on V_A_ estimates for fitness, we expected the highest relative fitness increase in the most stressful condition Hot-Dry and similar increases in single stress conditions Dry and Hot. High adaptive potential in females in Hot was reduced by a negative covariance with male fitness. We tested adaptation to the three conditions after 20 generations of experimental evolution and found that observed adaptation mainly matched our predictions. Given that body size is commonly used as a proxy for fitness, we also tested how this trait and its genetic variance (including non-additive genetic variance) were impacted by environmental stress. In both traits, variances were sex and condition dependent, but they differed in their variance composition, cross-sex and cross-environment genetic covariances, as well as in the environmental impact on V_A_.

## Introduction

Environmental changes pose a substantial risk of extinction to many organisms (Thomas et al. 2004; Parmesan 2006). Predicting whether a population is able to persist is therefore of crucial importance. Species may adapt via plastic or genetic changes. Plastic changes, e.g. physiological or behavioural adjustments, allow individuals to cope with stressful conditions (Charmantier et al. 2008). However, plastic responses are often costly and thus likely limited (Houle 1992; DeWitt et al. 1998; de Jong 2005; Valladares et al. 2007; Pfennig et al. 2010; Snell-Rood et al. 2010; Sokolova et al. 2012). It is also not clear whether they are sufficient to compensate negative effects of environmental changes (Duputié et al. 2015; Arnold et al. 2019). Instead, fast environmental shifts may require rapid adaptation by genetic evolution on short time scales. In this case, the standing genetic variation already present in the population is of particular importance (Kellermann et al. 2006; Bell and Gonzalez 2009). Adaptive evolution proceeds through different individual contributions to the next generation, i.e. differences in their fitness. Importantly, this variation in individual fitness has to be at least partly due to some underlying genetic variation in order to change the genetic composition of a population. Usually, only variance due to additive genetic effects (V_A_) is considered since additive effects are inherited from parents to offspring and determine the response to selection. V_A_ of relative fitness gives the expected increase of fitness in the next generation (Fisher 1930; Price 1972; Falconer and MacKay 1996). Thus, the existence of V_A_ in fitness is a prerequisite for adaptation and can be used as an overall estimate of a population’s capacity to adapt, or its evolvability (Houle 1992; Hansen et al. 2011; Shaw and Shaw 2014).

A common and intuitive expectation is that V_A_ in fitness should be low because selection depletes genetic variation in adapted populations. Accordingly, it was found that heritability (h^2^, proportion of V_A_ in total variance V_P_) was lower in traits more closely associated with fitness (Mousseau and Roff 1987; Kruuk et al. 2000; Merilä and Sheldon 2000; Teplitsky et al. 2009; Wheelwright et al. 2014). However, h^2^ is not ideal for estimating V_A_ in fitness and the evolutionary potential of a population (Hansen et al. 2011; Wheelwright et al. 2014; Morrissey and Bonnet 2019) since low h^2^ is often due to higher environmental variance (Schluter et al. 1991; Merilä and Sheldon 1999). Many populations may also not be at an evolutionary equilibrium (Shaw and Shaw 2014) in environments that have been recently changed by human impacts (Fugère and Hendry 2018). Estimates of V_A_ in fitness are scarce, especially in natural populations and tend to show large heterogeneity in the estimates of V_A_ for lifetime fitness (e.g., in vertebrate populations: Kruuk *et al*., 2000; Merilä & Sheldon, 2000; McCleery *et al*., 2004; McFarlane *et al*., 2014). Although it is predicted that some species might not have the evolutionary potential to adapt to shifting environmental conditions (Etterson and Shaw 2001), some studies found substantial amounts of V_A_ in fitness in wild populations. Similarly, laboratory studies have reported significant V_A_ for fitness related traits in *D. melanogaster* (Gardner et al. 2005; Fry 2008; Long et al. 2009).

Despite existing V_A_ in fitness traits, evolution to new conditions can be constrained by antagonistic pleiotropy if alleles influence several fitness components but with opposite effects. This leads to trade-offs since one component cannot be optimized without reducing the other, e.g. fecundity and life length (Roff 2000). Another example of trade-offs is sexual antagonism (Foerster et al. 2007; Bonduriansky and Chenoweth 2009; Delcourt et al. 2009; Kirkpatrick 2009; Poissant et al. 2010; Calsbeek et al. 2015; Connallon and Hall 2016). Fitness optima might often differ between males and females. However, sharing a great part of their genome constrains independent evolution and limits adaptation when selection in the two sexes is opposite to the genetic correlation. Such a constraint should be revealed by negative genetic correlations between male and female fitness. Similar constraints may appear when adaptation to certain environmental conditions (e.g., elevated temperature) trades-off with adaptation to other conditions. Environmental changes often include simultaneous changes in several variables. For short-term stress exposure, effects like cross-tolerance and hardening (i.e., resistance to one stress develops after exposure to another stress) were observed, and it is known that some stress responses rely on the same physiological mechanisms (Bubliy et al. 2012). Evolutionary adaptation to different stressors involving the same pathways may thus lead to correlated resistance to another stressor (Bubliy and Loeschcke 2005; Sikkink et al. 2015). However, examples of local adaptation suggest that selection may often favour different genotypes in different conditions (Hereford 2009). Immediately after exposure to an environmental change, genetic correlations between fitness in different conditions can inform us to what extent the genetic basis of fitness is shared between environments. If we observe negative genetic covariances between fitness in different conditions, it is likely that alleles providing fitness benefits in one condition become detrimental in another. Adaptation when both stress factors are experienced at the same time can then be limited.

Changes in environmental conditions are also well known to affect genetic variances (Sgrò and Hoffmann 1998; Hoffmann and Merilä 1999; Rowiński and Rogell 2017), and covariances (Simons and Roff 1996; Sgrò and Hoffmann 2004; Wood and Brodie 2015), including cross-sex genetic covariance (Delcourt et al. 2009; Poissant et al. 2010; Punzalan et al. 2014). This can have substantial implications for evolutionary potential (Wilson et al. 2006; Husby et al. 2011), since the environmental shift that imposes a risk of extinction can at the same time either increase V_A_ for fitness (Shaw and Shaw, 2014) or reduce the evolutionary potential to adapt to this new condition (Wood and Brodie 2016). To fully understand the impact of environmental change on a population’s persistence it is therefore essential to know how genetic variances change in different environments and to identify potential constraints.

The fundamental importance of V_A_ of fitness for predicting contemporary evolution (Shaw and Etterson 2012; Hendry et al. 2018) and recent statistical advances in quantitative genetics have fostered great interest in estimating the adaptive capacity of wild populations (Charmantier et al. 2014). Even more so with the recent progress in using genomic markers to infer genetic resemblance among individuals (Gienapp et al. 2017; Perrier et al. 2018). These advances open up new perspectives for applications of classical quantitative genetic and genomic tools in wild populations, addressing important questions regarding populations’ persistence under environmental change (Waldvogel et al. 2020). However, so far relatively little information exists about the predictive value of V_A_ for fitness over several generations. In our study, we addressed this question by combining classical quantitative genetics with experimental evolution in the model organism *Tribolium castaneum*. We used a two-generation half-sib/full-sib breeding design to estimate genetic variances of fitness traits in four different conditions (control, dry, hot and hot-dry) representing two often co-occurring stressors, heat and drought. We measured offspring number as estimate of fitness in the F1 generation, and body size in the F2 generation, as an additional trait, often use as fitness proxy. We evaluated adaptation to heat, drought and their combination after 20 generations of experimental evolution. Thus, our experimental setup allowed us to explore many different facets of adaptation and to ask: how the adaptive potential changes under stressful conditions, whether trade-offs between female and male fitness can constrain adaptation, and by testing for genotype-by-environment interactions (G x E,) to which extent resistance to different stressors shares a common genetic basis. Having obtained such estimates in the founder populations, we could then gain deeper insights into the process of adaptation. Linking experimental evolution and classical quantitative genetics proved to be a powerful approach to evaluate the predictive power of genetic variance for fitness, and obtain a better understanding of the observed adaptation after 20 generations in new environments.

## Material and Methods

### Animal rearing and stress treatments

We used the *Tribolium castaneum* Cro1 strain (Milutinović et al. 2013), an outbred lab strain collected from a wild population in 2010, kept at high population size (>10,000) and adapted to lab standard conditions (33°C, 70% relative humidity) for more than 30 generations. Beetles were kept in 24h darkness on organic wheat flour mixed with 10% organic baker’s yeast. We sterilized the flour and yeast by heating them for 12h at 80°C before use. We measured fitness as well as size in control (=standard) conditions and three stress treatments with increased temperature or/and decreased humidity. The conditions in the treatments were: Hot: 37°C and 70% relative humidity, Dry: 33°C and 30% r. h., Hot-Dry: 37°C and 30% r. h. In order to be able to estimate genetic variances, we applied a split-brood paternal half-sib breeding design. We produced 147 half-sib families by mating virgin males to three virgin females (Figure S1A). Half-as well as full-sib families were split across all conditions (Figure S1A). Male and female offspring (four females and two males per full-sib family and condition) were separated at the pupal stage and transferred to 10 mL tubes with 1 g of medium and remained there until they were used for the fitness assay eight weeks later.

### Fitness Assay and producing double first cousins

To estimate fitness, we mated each virgin male with two unrelated virgin females from the same condition in 15mL tube with 1g medium. The male was removed after 24h and females transferred into two separate tubes. Females were removed from the tubes after one week of egg laying, and 9g medium was added to provide food for the developing offspring. After five weeks the number of adult offspring was counted. While we conducted the matings for the fitness assay, we followed a specific crossing design and always crossed two pairs of full-sib families (Figure S1B). Individuals resulting from these crosses (F2) were double first cousins.

### Size

Body size was measured in the F2, i.e. in the offspring of beetles that were used for the fitness assay. To estimate body size, we used the centroid size of the abdominal segment IV as proxy for total size since it can be measured more accurately than dry weight in very small insects and shows a high correlation with body mass (Wickman and Karlsson 1989; Honěk 1993). We decided to use size of the abdominal segment because we could measure it with higher accuracy than the total size, which strongly depends on whether an individual is in perfectly stretched position. The size of other parts of the body show a high correlation with this segment (Supplemental Figure S3). Dead beetles were fixed with a double-faced Scotch tape dorsally on a microscope slide. Sex was determined based on the sex patches of males at the inside of the femur. After sexing, all legs of the beetles were removed. Slides with single specimens were placed under a Wild M8 Heerbrugg M8 dissection microscope with a transmitted light stand. Two further light sources from above were installed for better illumination. Images were captured with a 25x magnification using a Leica DFC495 digital camera connected to a PC running the Microscope imaging software LAS v4.6.2. File utility program “tpsUtil” was used to build tps files from the images and tpsdig2 for setting the landmarks (http://life.bio.sunysb.edu/ee/rohlf/software.html for program information). To estimate beetle size, four landmarks were set on the ventral part of the abdominal lV (https://figshare.com/articles/_Morphology_of_the_red_flour_beetle_Tribolium_castaneum_/759706/1). The centroid size was calculated using the free software Past 3.14 (for information see: http://folk.uio.no/ohammer/past/).

### Effects of condition on fitness and size

Statistical analysis was conducted in R version 3.4.2 (R Core Team 2017). We used linear mixed models as implemented in the Rpackage *lme4* 1.1-17 (Bates et al. 2015) to explore the effects of treatments on fitness and size. For fitness analysis we included treatment and batch (time of fitness assay) as fixed effects and mother identity as random effect to account for non-independence since some of our measured individuals were full-sibs. For analysing size we included batch (individuals that grow up at the same time), treatment and sex as well as their interaction as fixed effects and mother identity as random effect. Significance and confidence intervals were obtained using the Rpackages *lsmeans* 2.27-62 (Lenth 2016) and *lmertest* 3.0-1 (Kuznetsova et al. 2017).

### Adaptation

We used ten replicate lines per condition originating from the same ancestral population (same as before, Cro1) and let them adapt for 20 generations (Supplemental Figure S2). Each new generation was set up by randomly selecting 120 pupae and placing them into a new vial with 70g medium. One selection line in Dry became extinct. Adult beetles of generation 20 from all selection lines were transferred to control conditions, in which they stayed for one week to mate and lay eggs. After removal of the adults, we waited until their offspring had reached the pupal stage and separated males and females. These individuals (generation 21) developed completely in control conditions. When they had reached the adult stage, each virgin male was mated with a virgin female of the same selection line and their offspring was transferred to all four conditions in the egg stage, resulting in full-sib families split across all conditions (Figure S2). As soon as these offspring (generation 22) had reached the pupal stage, males and females were separated. To compare fitness of different selection lines and test for adaptation, a virgin male and a virgin female of the same selection line in the same condition, but from different families were mated and the number of adult offspring produced within four days of mating and egg laying was used as a fitness estimate. To test for adaptation, we compared whether offspring number of selection lines in their native condition was significantly higher compared to non-adapted control lines. First, we analysed each condition separately. We used linear mixed models including selection regime as fixed effect and lines and families nested within lines as random effects using the Rpackage *lme4* (Bates et al. 2015). We were further interested to test for correlated responses, i.e. whether adaptation to a certain stress treatment could increase fitness in another. For this we run one analysis using the complete data set. Selection regime, conditions, and their interaction were used as fixed effects, lines, families nested within lines, and line-treatment interaction as random effects. Reported effect sizes and standard errors were obtained from the summary output of the model. p-values and confidence intervals were computed with the Rpackages *lmertest* (Kuznetsova et al. 2017) and *lsmeans* (Lenth 2016). Statistical analyses were conducted in R (R Core Team 2017). Model diagnostic plots are shown in supplemental Figure S6 and the complete results in Table S2.

#### Quantitative genetic analyses

We estimated genetic variances, covariances and correlations using an animal model and restricted maximum likelihood estimation as implemented in Asreml version 3.0 (Gilmour et al. 2009). The animal model is a linear mixed effect model that uses all known relationships from a pedigree as random effect to partition observed variance into additive genetic variance and other sources of variance (Kruuk 2004). Non-additive genetic relationship matrices were created with the Rpackage *nadiv* 2.16.0.0 (Wolak 2012). All models were run in R by Asreml-R (Butler et al. 2009). All models reached convergence.

### Fitness

First, we ran a series of univariate models for each condition separately with offspring number as response variable, batch (samples where fitness assay was started on the same day) as fixed effect, additive genetic effects of females and additive genetic effects of males as random effects. Maternal identity (mother of female) was included as random effect to account for resemblance of full-sibs due to maternal, common environment or non-additive genetic effects. Significance of random effects was determined by likelihood ratio tests, testing whether excluding a certain random effect resulted in a significantly worse model. Reported P-values are one-tailed since we tested whether a certain variance is different from zero and variances cannot become negative (Wilson et al. 2010). Although count data, it can be analysed assuming a Gaussian distribution when distribution converges towards a normal distribution (de Villemereuil 2018). Distribution of offspring numbers are shown in Figure S4. Model diagnostic plots and more details are given in the Supplement (Figure S5). For estimating cross-sex genetic correlation we used the same univariate models but included a covariance between female and male additive genetic effects. Maternal effects were removed in subsequent analyses since they were very small and non-significant. Studies so far used bivariate models to investigate cross-sex genetic correlations (Brommer et al. 2007; Foerster et al. 2007; McFarlane et al. 2014; Punzalan et al. 2014; Wolak et al. 2018). In case of fitness this could be problematic, because the same observation (number of adult offspring resulting from a mating) would be used twice. Using univariate models including additive genetic effects of females and males avoids pseudoreplication. Significance of genetic correlations was determined by two-tailed likelihood ratio tests comparing a model including correlations to a model with correlations set to zero. In order to test for genotype-by-sex interaction (G x S), which indicates that a genotype is differently expressed in males and females, we compared the model to a constrained model with genetic correlation fixed to one. To examine whether additive genetic effects of females and males were different, we tested whether the unconstrained model was significantly better than a model with V _A_ of females and V_A_ of males forced to be equal within each condition.

For estimating cross-condition genetic covariances and correlations we used bivariate models with fitness in two conditions as response, batch as fixed and additive genetic effects of females and males as random effects. Since each individual was only measured in one condition, residual covariance was set to zero. Significance of correlation was assessed by comparing to a model with correlation fixed to zero using two-tailed likelihood ratio tests. Genetic correlations between traits measured in different environments that are significantly less than unity, indicate a G x E interactions (Kruuk 2004; Charmantier and Garant 2005). Comparison with a model with correlation fixed to one was used to test for G x E (Wilson et al. 2010). In all bivariate models, covariance between male effects was set to zero, because male effects were very small and not significant in some conditions and hence it was not meaningful to estimate covariances.

To allow comparisons across different conditions, we calculated h^2^ and CV_A_ (CV_A_ is the square root of V_A_ divided by the phenotypic mean of the trait, multiplied by 100) of fitness for males and females in each condition. I _A_ (V_A_ divided by trait mean squared, multiplied by 100 (Houle 1992)) was calculated as an estimate for the proportional change after one generation. Standard errors of CV_A_ and I_A_ were computed as described in Garcia-Gonzalez *et al*., (2012).

To assess whether genetic variances were significantly affected by environmental change, we used a multivariate model with fitness in each condition as separate response variable. Since fitness between conditions differed in mean and variance, we standardized data. We applied two different standardizations: 1) mean standardized (i.e. relative fitness) and 2) dividing by the standard deviation (sd) to have a variance of one. We then tested whether constraining the multivariate model and forcing V_A_ of standardized fitness to be equal in all conditions resulted in a significantly worse model. If the constrained models are worse, we can conclude that mean standardized V_A_ (I_A_ of absolute numbers), or sd standardized V_A_ (h^2^ of absolute numbers) are significantly influenced by the environment. Similarly, we tested whether genetic cross-condition correlations were different. We used a multivariate model and constrained all pairwise cross-condition correlations to be equal. We then tested if this model was significantly worse than an unconstrained model using a two-tailed likelihood-ratio test.

### Size

Our crossing design during the fitness assay (reciprocal crossing of two pairs of full-sibs, see Figure S2) allowed us to estimate non-additive genetic variance (V_D_) in the following generation (F2). In a paternal half-sib breeding design these effects contribute to resemblance among full-sibs and cannot be separated from maternal or common environmental effects. In contrast, double first cousins (DFC) share non-additive genetic effects without being confounded by maternal effects or a common environment. We partitioned the observed variance for size into additive (V_A_), maternal (V_M_), non-additive (V_D_), and residual variance V_R_ while controlling for batch effects. Batches represent individuals that grew up at the same time, and thus accounts for variations in the medium or lab temperature. Similar to the analysis of fitness data, we first analysed each condition separately using univariate animal models with batch as fixed and maternal, additive, and non-additive genetic effects as random effects. To get sex specific estimates, we analysed male and female size separately. Genetic correlations between sexes and between conditions were assessed by bivariate models. Significance of correlations, G x E, environmental effects on I_A_ and h^2^ of size were tested in the same way as we described before.

To demonstrate how responses are affected by covariances between the sexes we used estimated variances and covariances between male and female additive genetic effects of relative fitness to predict fitness increase after one generation. We applied the multivariate breeder’s equation *Δ z*=*Gβ* (Lande 1979), where *Δ z* is a vector of changes in the trait means, *G* the 2 × 2 genetic variance-covariance matrix for female and male relative fitness estimated in each condition, and *β* a vector of selection gradients. We used a vector of selection gradients that assumed equal selection on male and female fitness, β^T^=[1,1]. Total change in fitness is then the sum of fitness increase in females and in males. Alternatively, we may consider male additive genetic effects as indirect genetic effects on female reproductive output. Joint effects of direct and indirect selection on evolutionary changes of a single trait can be calculated as described in Bijma & Wade (2008) (equation 14):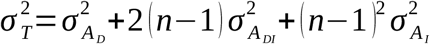, where 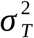 is the total heritable variance, 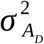 additive genetic variance for direct effects (female V), 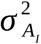 additive genetic variance for indirect effects (male V_A_), and *n* gives the number of interacting individuals (here, it equals two). In case of relative fitness, this total heritable variance is equal to the predicted relative increase in fitness (Fisher 1930). Thus, considering male and female fitness as two separate traits and applying the multivariate breeder’s equation, or treating additive genetic effects of males as indirect genetic effects on female fitness is equivalent.

## Results

### Effect of treatment

It was previously shown that all treatments had a highly significant effect on offspring number (Koch and Guillaume 2020) with the effect of heat being stronger and the lowest offspring number when heat and drought were combined. (Figure 1A). Treatments also showed a significant effect on size (F_3,7710_= 76.30, p < 2.20E-16, Table S1) and size decreased in the stressful conditions. Similar to offspring number, the effect of drought was smaller. Combining both stressors did not result in an additional decrease in size. In contrast, we observed the lowest size in Hot and higher sizes in Hot-Dry (Figure 2A). The difference between males and females was significant (F_1,7759_ = 518.19, p < 2.20E-16) with females being larger. We also detected a significant sex-by-condition interaction (F_3,7759_ = 7.27, p = 8.16E-04) indicating that males were more sensitive to stressful conditions (Figure 2A).

**Figure 1:**
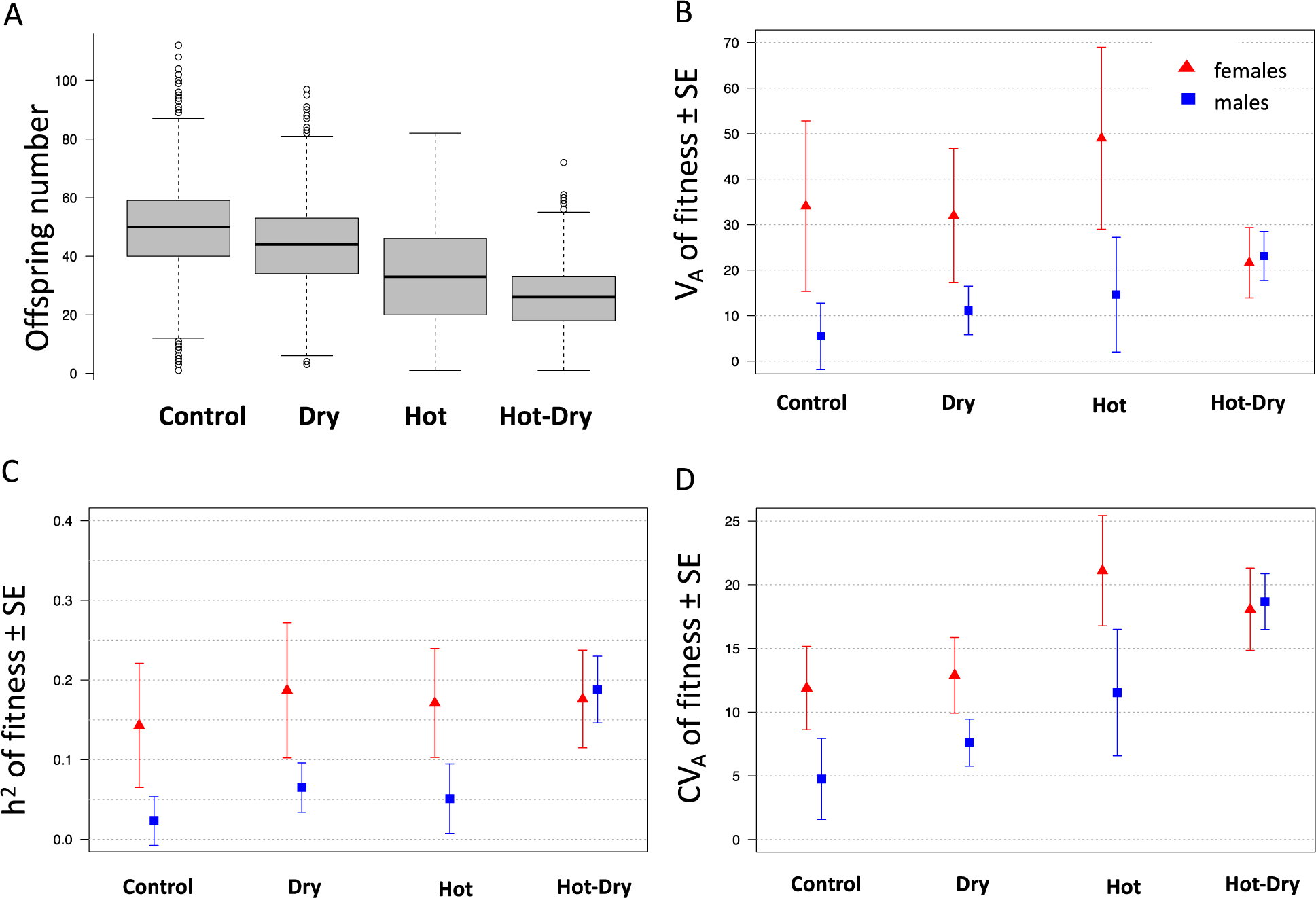
Fitness (= number of adult offspring) **(A)** and estimates for its additive genetic variance (V_A_) **(B)**, heritability (h^2^) **(C)**, and coefficient of V_A_ (CV_A_), which is the square root of V_A_ divided by the phenotypic trait mean, multiplied by 100 **(D)** in female and male flour beetles (*Tribolium castaneum*) in four different environmental conditions.

**Figure 2:**
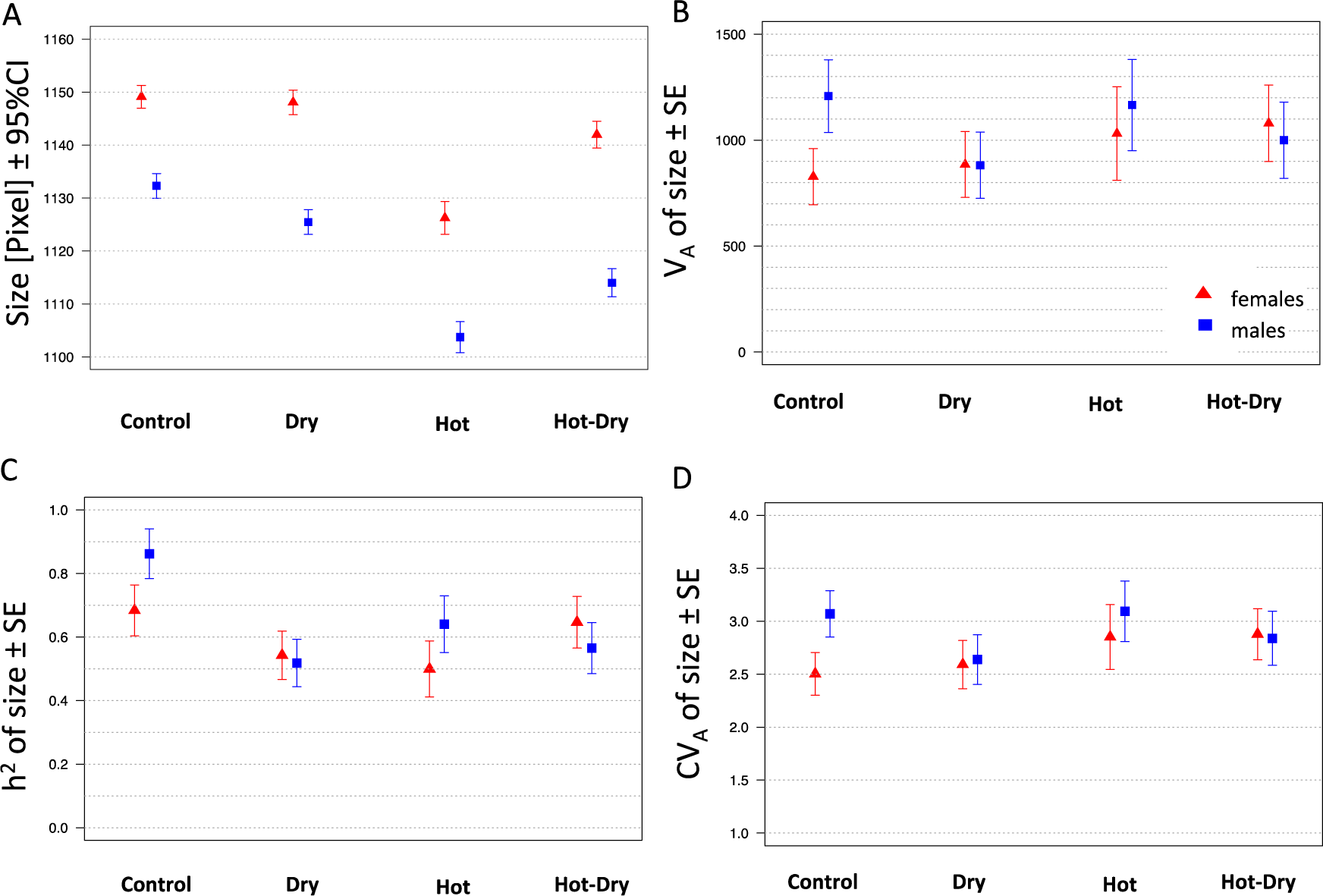
Body size (centroid size of abdominal segment IV) **(A)** and estimates for its additive genetic variance (V_A_) **(B)**, heritability (h^2^) **(C)**, and coefficient of V_A_ (CV_A_) **(D)** in female and male flour beetles (*Tribolium castaneum*) in four different environmental conditions.

### Genetic variances and covariances

#### Fitness

V_M_, which consisted of variance due to maternal effects, common environment and non-additive effects, was not significant in any condition. We found significant V_A_ for female fitness in all conditions (Table 1). V_A_ for males was generally smaller (P = 8.10E-03) and not significantly different from zero in Control and Hot. A remarkable exception was Hot-Dry, where male V_A_ was increased and of the same magnitude as female V_A_ (Figure 3). V_A_ in females was similar in Control and Dry but increased in Hot and was reduced in Hot-Dry (Figure 1B). Since residual variances changed simultaneously with V_A_ (Table 1), h^2^ for females was similar in all stress conditions (Figure 1C). When we compared genetic variances using standardized data, we found h^2^ of males (P = 2.75E-03), but not females (P = 0.55) to differ significantly between conditions. Similarly, I _A_ was affected by conditions in males (P = 8.87E-05), but not in females (P = 0.07). When we tested a model constraining V _A_ of females and males to be the same in all conditions, we found it to be significantly worse (P = 2.46E-08) than an unconstrained model, thus showing that the total amount of heritable genetic variance (V_A_ of females and males) was different between treatments.

**Table 1:** Genetic variances (V_A_: additive genetic variance; V_M_: maternal variance; V_R_: residual variance), heritability (h^2^), coefficient of additive genetic variance (CV_A_) of offspring number and cross-sex additive genetic covariances (COV_A_) and correlations (r_A_) in different environmental conditions. Estimates for genetic variances were obtained from a univariate animal model including additive genetic effects of males and females as random effects. For COV_A_ and r_A_ univariate animal models were used with covariance between female and additive genetic effects. N gives the number of reproducing females used for the analysis. All significant results are in bold.

|  | Condition |  |  |  |
| --- | --- | --- | --- | --- |
|  | Control | Dry | Hot | Hot-Dry |
| **N** | 1514 | 1603 | 1005 | 1396 |
| **mean (SE)** | 49.09 (0.4) | 43.88 (0.36) | 33.18 (0.55) | 25.73 (0.31) |
| **$V_A$ females (SE)** | **34.06 (18.75)** | **32.00 (14.71)** | **48.99 (20.02)** | **21.62 (7.71)** |
| **$h^2$ females (SE)** | **0.14 (0.08)** | **0.19 (0.09)** | **0.17 (0.07)** | **0.18 (0.06)** |
| **$CV_A$ females (SE)** | **11.89 (3.27)** | **12.89 (2.97)** | **21.10 (4.32)** | **18.07 (3.23)** |
| **$V_A$ males (SE)** | 5.44 (7.29) | **11.14 (5.35)** | 14.62 (12.59) | **23.09 (5.40)** |
| **$h^2$ male (SE)** | 0.02 (0.03) | **0.07 (0.03)** | 0.05 (0.04) | **0.19 (0.04)** |
| **$CV_A$ males (SE)** | 4.75 (3.18) | **7.61 (1.83)** | 11.53 (4.97) | **18.68 (2.20)** |
| **$V_M$ (SE)** | 5.18 (10.30) | 4.90 (7.55) | 2.26E-07 (5.03E-0.5) | 2.76E-05 (2.21E-06) |
| **$V_R$ (SE)** | 193.39 (13.65) | 122.88 (9.79) | 222.27 (20.00) | 78.35 (6.27) |
| **$V_P$ (SE)** | 238.22 (8.88) | 170.91 (6.33) | 235.88 (13.10) | 123.05 (4.96) |
| **$COV_{A \text{ female,male}}$ (SE)** | **25.73 (6.29)** | **11.75 (4.92)** | -9.24 (11.02) | **11.54 (4.21)** |
| **$r_{A \text{ female,male}}$ (SE)** | **1.55 (0.66)** | **0.54 (0.22)** | -0.36 (0.45) | **0.52 (0.18)** |

**Figure 3:**
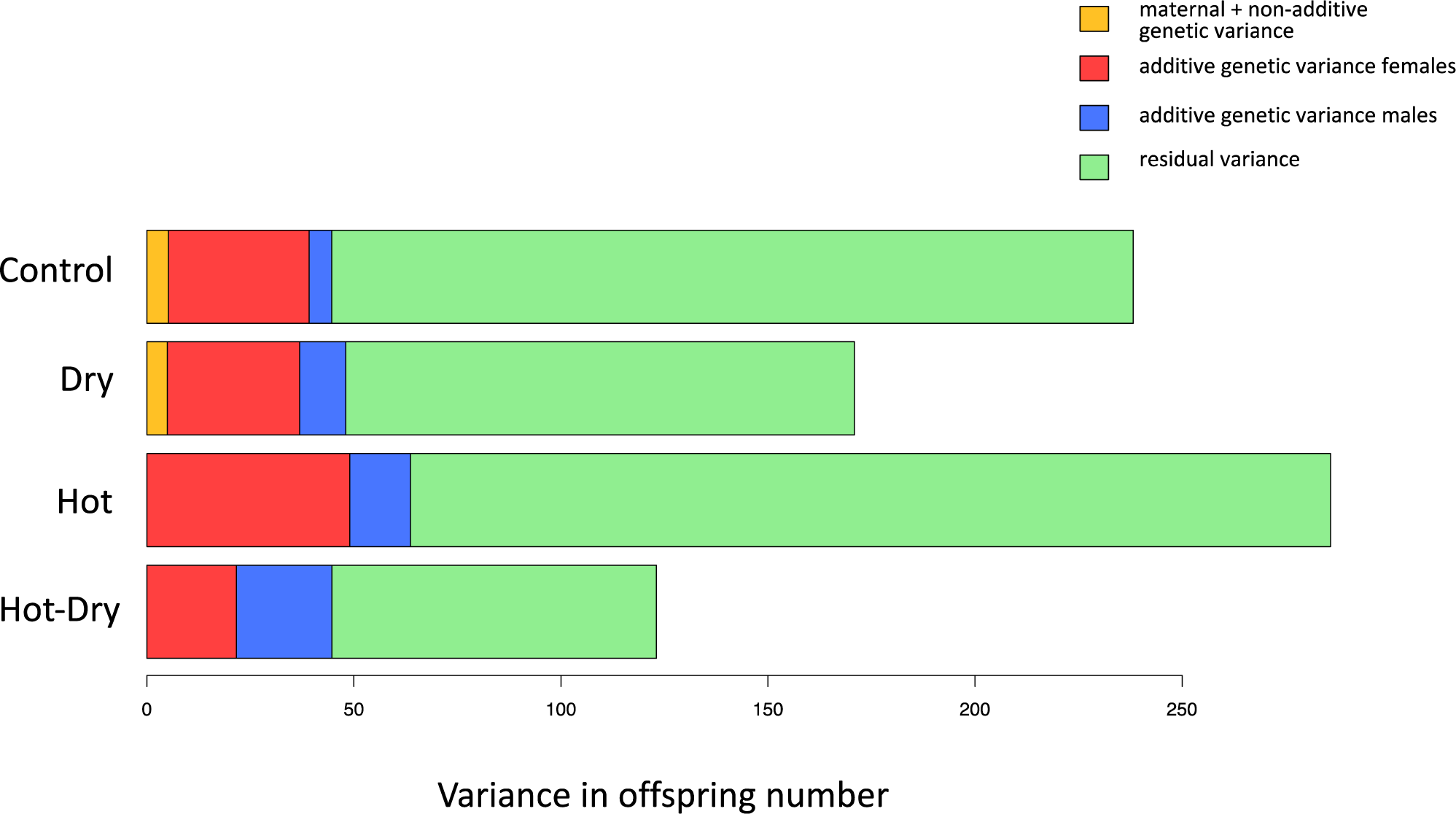
Variance components of offspring number of *Tribolium castaneum* in four different environmental conditions estimated using univariate animal models.

Genetic correlations between fitness in different conditions were always positive (Figure 4) and significant. However, we found significant differences between the different cross-condition correlations (P = 0.05). Correlations were slightly lower when the conditions differed in temperature and humidity (Control–Hot-Dry, Dry-Hot) (Figure 4). We found significant G x E between Control and Dry, Control and Hot-Dry as well as Dry and Hot.

**Figure 4:**
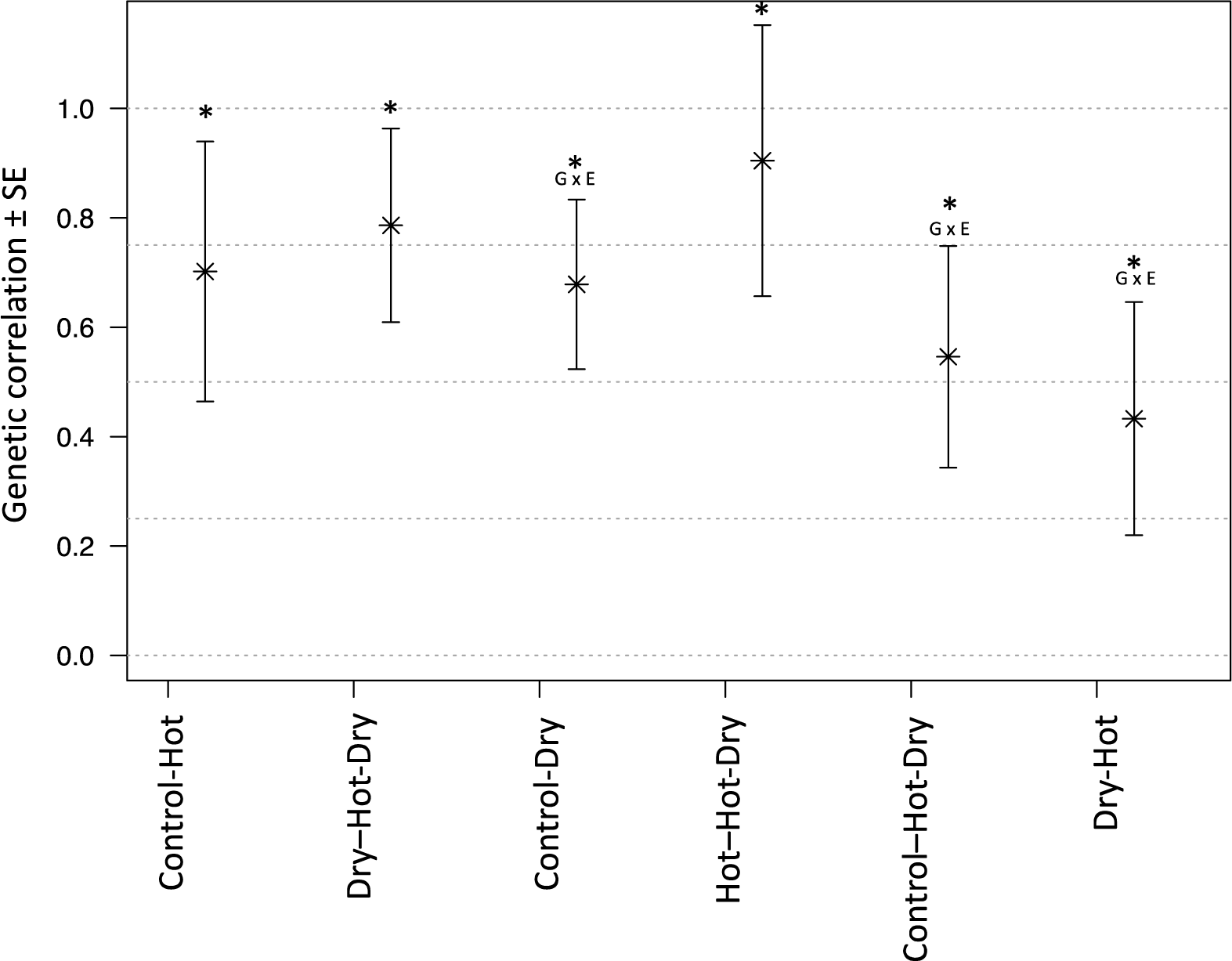
Pairwise cross-environment genetic correlations of fitness in *Tribolium castaneum* estimated using bivariate animal models. * show significant correlations, i.e. significantly different from zero. G x E (= genotype by environment interactions) indicate correlations significantly different from one. Control: 33°C, 70% relative humidity (r.h.); Dry: 33°C, 30% r.h.; Hot: 37°C, 70% r.h.; Hot-Dry: 37°C, 30% r.h.

Genetic correlations in fitness between females and males (Figure 5) were significant in Control (P = 1.02E-06), Dry (P = 1.34E-02), and Hot-Dry (P = 4.47E-04), but not in Hot (P = 0.10). The correlation was highest in Control (Table 1, Figure 5). It decreased in the stress treatments and became even negative in Hot. In all conditions except Control, we found that the correlation was significantly different from one (Dry: P = 1.95E-03; Hot: P = 5.78E-03; Hot-Dry: P = 1.26E-04), indicating G x S, i.e. that genetic basis of fitness is different in the sexes. I _A_, the expected evolutionary change as percentage of the mean, was highest in Hot and smallest in Control (females/males ± SE: Control: 1.59 ± 0.49/0.30 ± 0.29; Dry: 2.05 ± 0.50/0.63 ± 0.27; Hot: 4.32 ± 1.81/1.25 ± 1.14; Hot-Dry: 3.20 ± 1.14/ 3.51 ± 8.1). Based on this, we should observe adaptation in all stress treatments with the largest relative fitness increase in Hot. However, given the negative correlation between male and female additive genetic effects (Figure 5, Table 1), the evolutionary response may be constrained and less than predicted. Taking female – male genetic correlations into account (see Methods) gave us an estimated increase of mean fitness of 3.90% in Dry, 3.89% in Hot, and 10.20% in Hot-Dry per generation. Assuming V_A_ remains constant, the total change of mean fitness after 19 generations (generation 20) would be 106.81% in Dry, 106.57% in Hot and 532.54% in Hot-Dry.

**Figure 5:**
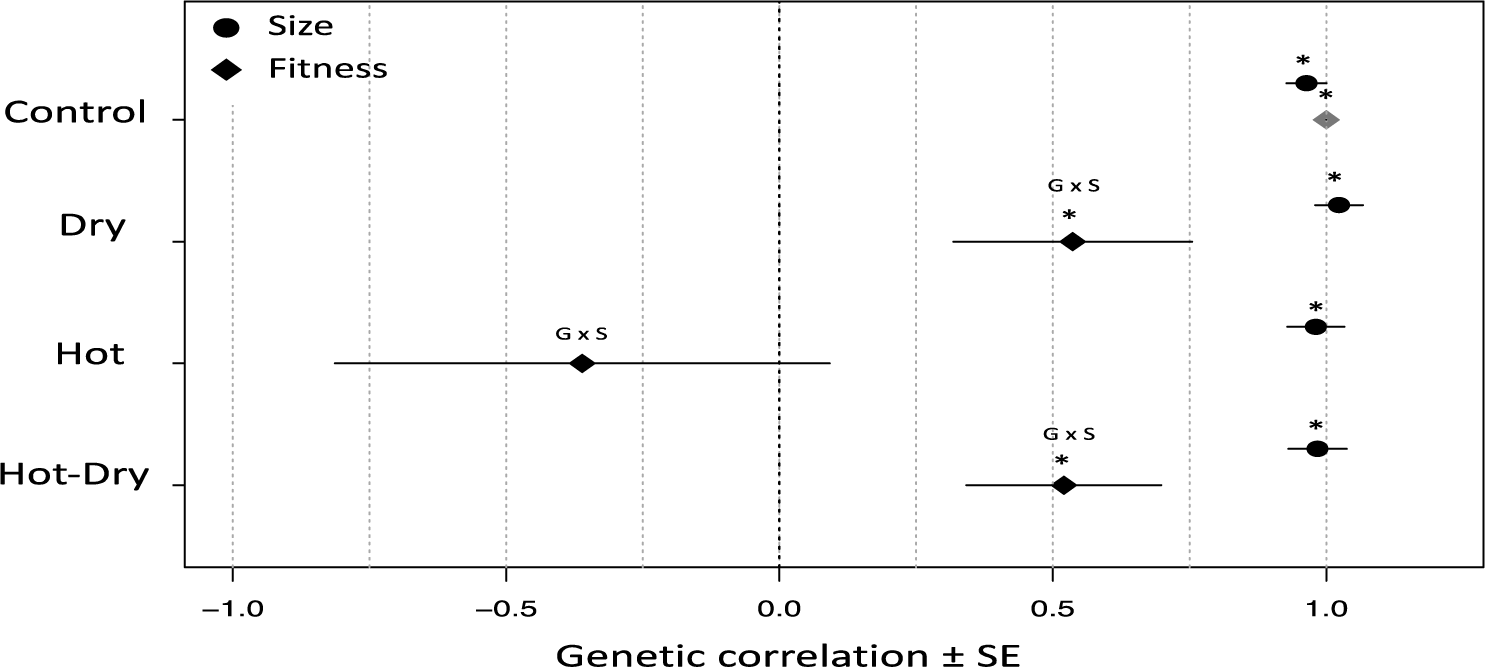
Cross-sex genetic correlations for size and fitness in *Tribolium castaneum* under different environmental conditions. * indicates a significant correlation. G x S (= genotype by sex interaction) indicates that correlation is significantly different from one. Estimate for genetic correlation of fitness in Control was bounded at one and no SE was available. Using models with unconstrained variances and covariances yielded an estimate of 1.55 ± 0.66. V_A_ of males was small in Control, which impedes precise estimation of cross-sex genetic correlation.

#### Size

V_A_ in size was highly significant in both sexes and in all conditions (P < 0.001). The results for V_D_ were less clear. In Control and Hot-Dry estimates for V_D_ were extremely small and estimates bounded at zero, indicating that non-additive effects contributed little to observed variation (Figure 6). In Dry and Hot, V_D_ was not significant, but variance estimates were high when it was included in the model (Table 2, Figure 6). A model without non-additive effects resulted in much higher estimates of V_A_ in Dry and Hot (Supplemental Table S3). In most cases, P-values for V_D_ were far from significance (P > 0.2), but for females in Dry and in Hot P-values were lower although still not significant (P = 0.13 and P = 0.10). When we combined males and females and added sex as fixed effect in the model, we obtained a P-value of 0.075 in Hot. Genetic correlations for size across conditions were always positive (Supplemental Figure S7), but no clear pattern emerged. We found significant G x E in female size between Control-Dry (P = 2.78E-03). In male size, G x E was significant between Control-Dry (P = 6.39E-03), Dry-Hot (P = 5.39E-03), Dry–Hot-Dry (P = 0.04), and Hot–Hot-Dry (P = 3.93E-03). Genetic correlations between female and male size were close to one (Table 2, Figure 5) in all conditions, suggesting that size cannot evolve independently in both sexes. I_A_ of female and male size was not significantly different (P = 0.08) considering all conditions. Although differences in Control seemed to be substantial (Figure 4B-D), they were not significant (h^2^: P = 0.26; I_A_: P = 0.06). We found that environmental change did not influence I_A_ and h^2^ of female size (P = 0.13; P = 0.14) nor male I_A_ (P = 0.13), but had a significant effect on male h^2^ (P = 0.02).

**Table 2:** Body size genetic variances (V_A_: additive genetic variance; V_D_: dominance variance; V_M_: maternal variance; V_R_: residual variance), heritability (h^2^), coefficient of additive genetic variance (CV_A_) in female and male flour beetles and cross-sex additive genetic covariances (COV_A_) and correlations (r_A_) in different environmental conditions. Estimates for genetic variances were obtained from univariate animal models for each sex separately. For COV_A_ and r_A_ bivariate animal models were used. All significant results are in bold.

|  |  | Condition |  |  |  |  |  |  |  |
| --- | --- | --- | --- | --- | --- | --- | --- | --- | --- |
|  |  | Control |  | Dry |  | Hot |  | Hot-Dry |  |
| Females | N | 1020 |  | 1181 |  | 837 |  | 960 |  |
|  | mean (SE) | 1149.13 | (1.10) | 1148.08 | (1.18) | 1126.25 | (1.57) | 1141.96 | (1.29) |
| | $V_A$ (SE) | **751.79** | **(201.63)** | **560.98** | **(305.17)** | **603.78** | **(446.35)** | **1017.45** | **(291.52)** |
| | $V_D$ (SE) | 6.63E-04 | (1.73E-04) | 379.526 | (782.07) | 333.554 | (1342.287) | 8.670E-04 | (2.21E-04) |
| | $V_M$ (SE) | 29.61 | (63.13) | 38.57 | (119.09) | 107.14 | (210.86) | 25.58 | (96.18) |
| | $V_R$ (SE) | 417.11 | (108.70) | 613.14 | (490.95) | 973.23 | (854.49) | 618.81 | (157.41) |
| | $V_P$ (SE) | 1198.52 | (72.27) | 1592.19 | (82.03) | 2017.74 | (117.05) | 1661.84 | (100.31) |
| | $h^2$ (SE) | **0.63** | **(0.14)** | **0.35** | **(0.18)** | **0.30** | **(0.21)** | **0.61** | **(0.15)** |
| | $CV_A$ (SE) | **2.39** | **(0.32)** | **2.06** | **(0.56)** | **2.18** | **(0.81)** | **2.79** | **(0.40)** |
| Males | N | 1008 |  | 1183 |  | 834 |  | 962 |  |
|  | mean (SE) | 1132.31 | (1.18) | 1125.47 | (1.19) | 1103.73 | (1.50) | 1114.01 | (1.34) |
| | $V_A$ (SE) | **969.79** | **(406.67)** | **632.66** | **(335.68)** | **954.83** | **(479.70)** | **999.70** | **(179.66)** |
| | $V_D$ (SE) | 369.18 | (567.38) | 404.65 | (539.19) | 346.49 | (748.68) | 4.94E-03 | (7.90E-04) |
| | $V_M$ (SE) | 6.59E-06 | (5.07E-05) | 4.88E-04 | (2.18E-04) | 6.54E-04 | (5.00E-04) | 3.14E-04 | (5.02E-05) |
| | $V_R$ (SE) | 35.85 | (275.83) | 634.64 | (282.89) | 496.90 | (380.49) | 769.42 | (123.08) |
| | $V_P$ (SE) | 1374.81 | (293.93) | 1672.04 | (87.39) | 798.22 | (118.96) | 1769 | (98.50) |
| | $h^2$ (SE) | **0.71** | **(0.25)** | **0.38** | **(0.19)** | **0.53** | **(0.25)** | **0.57** | **(0.08)** |
| | $CV_A$ (SE) | **2.75** | **(0.58)** | **2.23** | **(0.59)** | **2.80** | **(0.70)** | **2.84** | **(0.26)** |
| $COV_{A \text{ female,male}}$ (SE) | | **955.25** | **(120.93)** | **1002.06** | **(128.10)** | **1140.11** | **(175.33)** | **976.20** | **(137.10)** |
| $r_{A \text{ female,male}}$ (SE) | | **0.96** | **(0.04)** | **1.02** | **(0.04)** | **0.98** | **(0.05)** | **0.98** | **(0.05)** |

**Figure 6:**
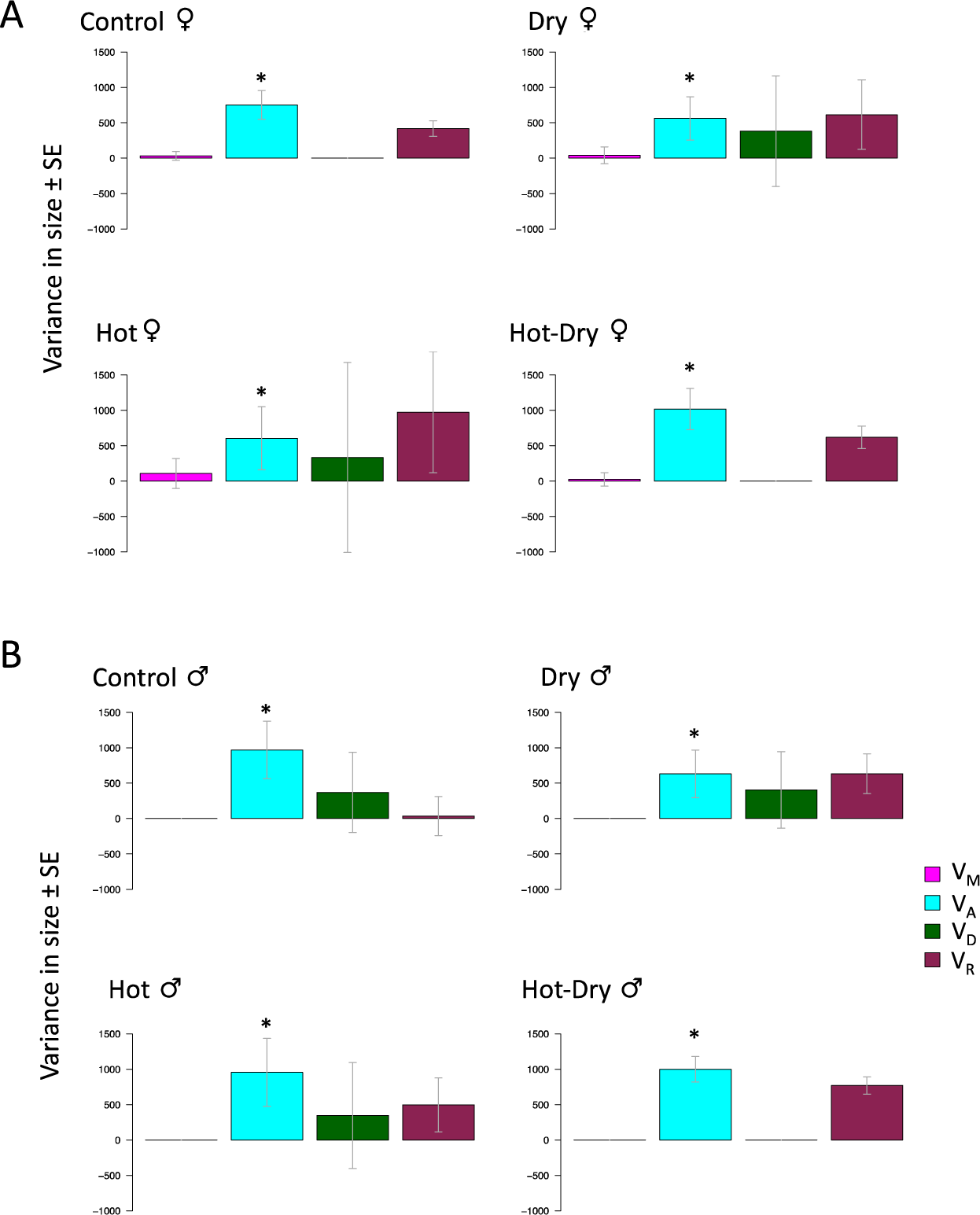
Variance components of size in female **(A)** and male **(B)** flour beetles (*Tribolium castaneum*) under four environmental conditions: V_M_: maternal variance, V_A_: additive genetic variance, V_D_: non-additive genetic variance, V_R_: residual variance. * indicates a significant variance component.

### Adaptation after 20 generations and asymmetric correlated responses

We found significant adaptation to all conditions after 20 generations, shown by significant effects of selection regime on offspring number (Dry: F_1,16_ = 10.43, p = 0.005; Hot: F_1,18_ = 4.80, p = 0.042; Hot-Dry: F_1,18_ = 14.78, p = 0.001, Table S2). In all treatments, the native selection lines produced significantly more offspring than non-adapted control lines (Figure 7). The largest difference and most significant fitness increase was observed in the most stressful condition Hot-Dry. In contrast to the three stress treatments, we did not find any differences between lines from different selection regimes in the ancestral control condition (Figure 7), showing that adaptation to stress treatments did not come at a cost of reduced fitness in control conditions. Interestingly, drought selection resulted in higher heat resistance. Dry lines showed a significantly higher offspring number in Hot and Hot-Dry (estimated increase in offspring number relative to Control lines: 12.10 ± 2.23, p = 3.64E-07 in Hot; 12.18 ± 2.18, p = 4.20E-07 in Hot-Dry). In Hot, they performed even better than the native Hot-lines (mean offspring number in Hot [95% CI]: Control-lines: 24.59 [20.71, 28.47]; Hot-lines: 29.94 [26.11, 33.77]; Dry-lines: 35.69 [31.77, 39.61]; Hot-Dry-lines: 36.91 [33.14, 40.68]). We did not observe such a correlated response in the Hot-lines (Figure 7). Their offspring number in Dry was not different from those of Control-Lines. Lines in Hot-Dry that adapted to a combination of heat and drought showed an increased fitness in both single stressor treatments (in Dry: 4.45 ± 2.10, p = 0.04; in Hot: 11.66 ± 2.17, p = 1.20E-04). Despite high genetic correlation in fitness between Hot and Hot-Dry estimated in the first generation, we found no correlated response of Hot-lines to Hot-Dry conditions. Their offspring number was not different from non-adapted Control-Lines (Hot-lines: 23.14 [19.43, 26.84 CI]; Control-lines: 23.03 [19.30, 26.76 CI]).

**Figure 7:**
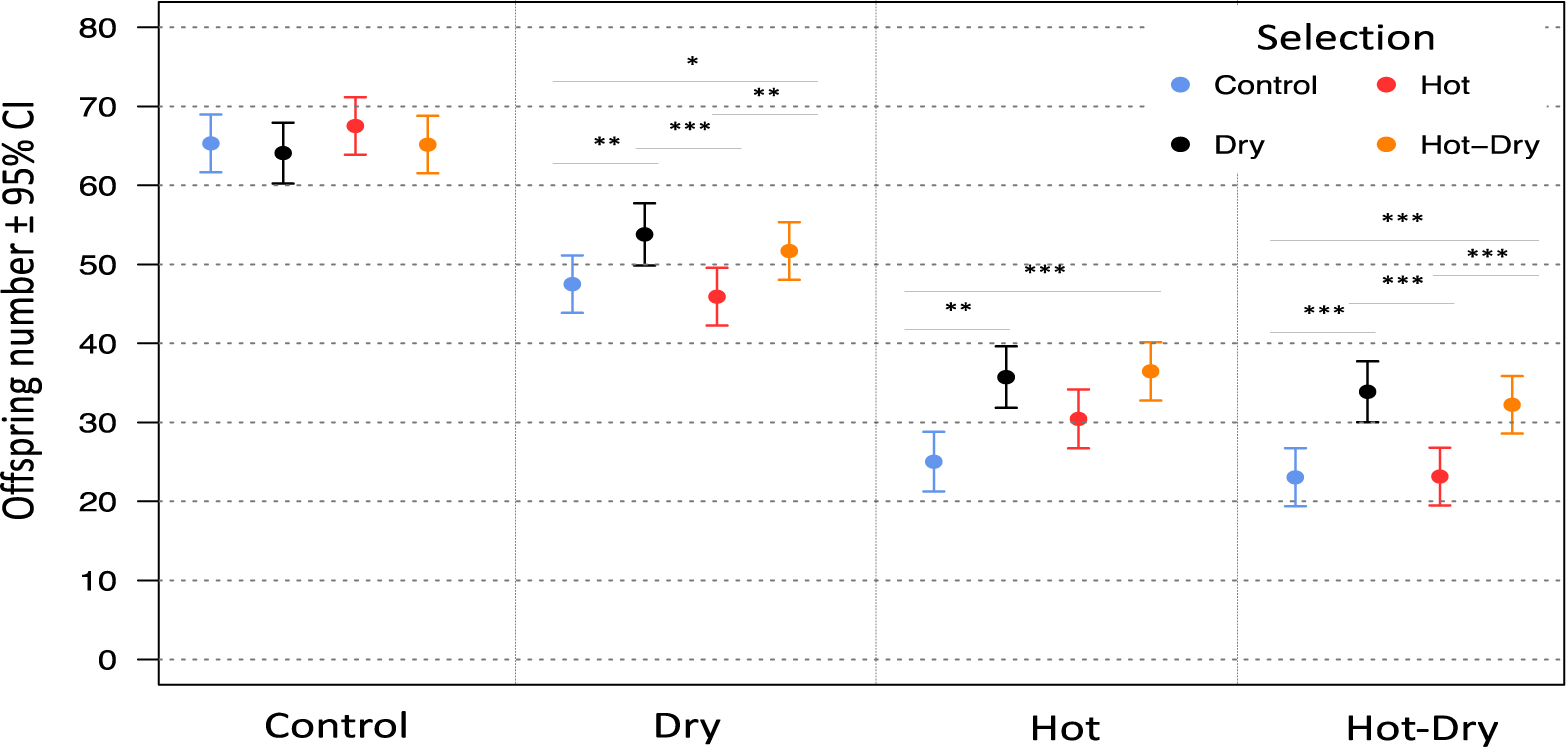
Mean offspring number per female of lines from different selection regimes under different environmental conditions. Significant differences between selection regimes within the same condition (Tukey’s HSD post hoc test, p-values adjusted for multiple comparisons) are indicated by *P<0.05, **P<0.01, ***P<0.001.

The observed changes in fitness after experimental evolution were much lower than the predicted total changes. Observed fitness increases relative to Control-lines were: Dry: 16.43 ± 3.24 %, Hot: 22.37 ± 4.34 % and 46.29 ± 2.17 % in Hot-Dry. Nonetheless, in agreement with our predictions we observed the strongest increase in Hot-Dry and similar increases in Dry and Hot.

## Discussion

Consequences of environmental change on persistence of populations strongly depend on their ability to adapt either by plastic or genetic changes. The stress conditions we applied, had a strong impact on fitness. Offspring number in Hot-Dry was reduced to ca. 50% of control level thus clearly showing that plasticity was not able to fully compensate for the negative effects of environmental change. We found significant V_A_ in fitness in all conditions indicating potential to adapt by genetic changes with no evidence that environmental changes lowered the adaptive potential. In contrary, I_A_ of female and male fitness increased in the stress treatments, which should facilitate adaptation. Accordingly, we found significant adaptation to all conditions after 20 generations of experimental evolution. We also did not find strong evidence of genetic constraints on adaptation between stressors or sexes as most genetic correlations were positive (but not in Hot, see Figure 5). Taking account of both male and female variances and covariance in fitness allowed us to make better predictions of fitness increase. All in all, our study shows that although it is not possible to predict the precise fitness increase over this period of 20 generations, we can make qualitative predictions about adaptation based on V_A_ estimates. We can examine whether a population is likely to adapt and make relative comparisons between stress treatments, i.e. identify those where we expect the largest relative increase in fitness. We thus show that existing quantitative genetic tools are informative over time scales beyond single generation responses. However, reliability of predictions may require a fully integrative approach including potential genetic covariances between the sexes as well as a careful choice of the traits used as fitness estimate.

### Evolutionary potential and adaptation

V_A_ of female fitness was higher than V_A_ of male fitness in all conditions except Hot-Dry. Egg production is likely to be costly and reduced when females are stressed due to a resource allocation trade-off. It is therefore not surprising that variation in reproductive output could be mainly explained by genetic differences among females, whereas genetic variation among males had only a minor influence. However, this changed when humidity was reduced. In Dry conditions, V_A_ for males became significant (but was still much smaller compared to females) and in Hot-Dry, V_A_ in both sexes were similar. It was observed that the condition of *Tribolium* males could have a significant influence on the reproductive output of their mating partner. Starvation of males reduced insemination success (Lewis et al. 2012), the number of eggs laid, and the proportion of unfertilized eggs (Sbilordo et al. 2011). It seems plausible that desiccation might have similar effects. Producing ejaculate may be costly and decreased under dry conditions to reduce water loss. Genetic differences in male drought resistance would then translate into observed offspring variation and result in a higher male influence on female reproductive output. Although male V_A_ was small in ancestral conditions, evolution of males seemed to play an important role in adapting to Hot-Dry. In contrast to the between-sex differences in fitness V_A_, we did not find clear differences in V_A_ between sexes for body size. The largest difference was in Control (Figure 2), with a lower female V_A_. It is likely that female size had been under strong positive selection since it is associated with fecundity (Honěk 1993), which might explain the observed reduction. Additionally, while we observed an increase in V_A_ for male fitness in the Dry treatment, V_A_ for male size was reduced in Dry. A drought effect was thus also detectable for size: When we explored the effects of temperature and humidity on size, we found a significant (although small) sex-treatment-interaction indicating that male size was more sensitive to drought than female size. G x E for male size occurred when conditions differed in humidity.

Positive cross-condition genetic correlations indicated an absence of an evolutionary trade-off between drought and heat adaptation. Consistently, we did not find any selection line performing worse than control lines in any condition after experimental evolution. It was previously shown that the treatments induce substantial changes in gene expression (Koch and Guillaume 2020). It might then be that selection in the stress treatments is on genes that are not expressed in control and thus allele frequency changes do not sufficiently affect fitness under control conditions. Because positive genetic correlations indicate a similar genetic basis for fitness under different conditions, adapting to one condition should result in a correlated response and increased fitness in other conditions. Accordingly, we found that drought adaptation improved heat resistance (Figure 7). However, this correlated response was asymmetric since we did not observe an effect of heat adaptation on drought resistance. A possible explanation is that selection in dry and hot conditions shifted allele frequencies of genes with different pleiotropic effects (Bohren et al. 1966). Alternatively, the pleiotropic degree of a gene might be environment dependent (Barrett et al. 2009). Interestingly, we found that genetic correlations in generation one were lower when the conditions differed in both temperature and humidity, which is the case in Control–Hot-Dry and Dry-Hot (Figure 4) suggesting that the genetic basis for adaptation to heat and drought might be slightly different.

We found the highest positive cross-sex genetic correlation in fitness in Control with lower estimates in the treatments and a negative correlation in Hot. Several studies in the wild (Brommer et al. 2007; Foerster et al. 2007; Mokkonen et al. 2011) as well as in laboratory populations (Delcourt et al. 2009; Punzalan et al. 2014) reported negative genetic correlations between female and male fitness. It is not well understood how cross-sex correlations are influenced by the environment and contrasting predictions have been made. For instance, it was argued that in a population far from its optimum after an environmental shift, females and males might experience similar directional selection leading to high a positive correlation in their fitness (Berger et al. 2014; Connallon and Hall 2016; Wolak et al. 2018). In contrast, genetic covariances between sexes have been predicted to be less negative in ancestral conditions (Delcourt et al. 2009), because selection on the long-term should favour alleles providing fitness benefits to both (Collet et al. 2016). This last prediction is consistent with our observations.

Taking the male V_A_ and the covariances between female and male fitness into account led us to predict the highest relative increase in fitness in Hot-Dry and smaller but similar increases in Dry and Hot. Ignoring male and cross-sex effects when studying the adaptive capacity of a population can thus lead to misleading predictions. The results of the fitness assay after 20 generations mainly matched our predictions but increases in fitness were much lower than our estimated upper limit. However, our predictions are based on the assumption of constant V_A_ and consequently exponentially increasing fitness (Falconer and MacKay 1996). Trade-offs between different fitness components are expected to occur because physiological limits, e.g. for egg laying rate, exist and should prevent an infinite fitness increase. It is also important to note that the conditions during the fitness assay were not exactly the same as during evolution. To measure the offspring number per female. each female was kept individually in egg-laying tubes. In contrast, all individuals of the same line were kept in one vial during experimental evolution. Effects of density and competition that might have influenced adaptation were not captured in our fitness assay. Our results after 19 generations might be further influenced by evolution of control lines that were used as reference representing the ancestral non-evolved stage. Significant V_A_ under control conditions in addition to high and positive genetic correlations between fitness in control and stress conditions suggest a fitness increase in control lines over time and correlated responses in the treatments. This might have led to an underestimation of the true fitness increase of adapted selection lines. Interestingly, we found no difference between lines from different selection regimes in Control. A reason might be that genes responsible for fitness in the treatments are not expressed under control conditions. Evolutionary changes in these genes that occurred during adaptation to the treatments would not show any effect on fitness under control conditions leading to cryptic genetic variation. In control lines these genes were not exposed to selection and consequently control lines showed a lower fitness compared to adapted selection line when exposed to the treatments.

### Variance components of fitness and body size

Both traits differed markedly in the proportion of different variance components and in the environmental effect on genetic variances. h^2^ was much higher in size than in fitness. This is a common finding in many studies (Roff and Mousseau 1987; Kruuk et al. 2000; McCleery et al. 2004; Teplitsky et al. 2009). It had been initially interpreted as evidence for strong selection depleting V_A_ in fitness related traits. However, a high V_A_ can be concealed in h^2^ by a simultaneous increase of the total variance (Houle 1992; Merilä and Sheldon 1999; Hansen et al. 2011; Wheelwright et al. 2014). Given the highly polygenic nature of fitness, it was even argued that fitness might show higher V_A_ (Merilä and Sheldon 1999), since it represents a larger mutational target.

Furthermore, fitness is a composite character with a high number of morphological, physiological and behavioral traits contributing to it, each of them with some underlying genetic variance. However, each contributing trait may increase the influence of environmental variation, leading to a higher total variance in fitness (Price and Schulter 1991) and thus a lower h^2^. According to those previous considerations, we found much higher estimates of mean-scaled V_A_ (CV_A_, I_A_) in offspring number than in body size and a lower h^2^ in fitness. The lower h^2^ was mainly due to a higher proportion of environmental variance (V_R_) when compared to body size. Contrary to body size, we could not directly estimate V_D_ for fitness because we could not disentangle V_D_ from V_M_ and common environmental effects in the F1 with our half-sib/full-sib breeding design. However, if V_D_ were present, then it should be included in V_M_, because it contributes to full-sib resemblance via a shared mother. Comparative studies investigating the relative amount of V_A_ and V_D_ suggested that proportion of V_D_ can be substantial and of the same magnitude as V_A_ (Crnokrak and Roff 1995; Wolak and Keller 2014). An increased proportion of V_D_ is expected in populations under strong selection, since V_D_ is not affected by natural selection (Crnokrak and Roff 1995; Merilä and Sheldon 1999; Roff and Emerson 2006; Sztepanacz and Blows 2015), or with increased inbreeding (Falconer and Makay, 1996). In our experiment, V_M_ for fitness was much lower than V_A_, even close to zero in Hot and Hot-Dry (Figure 3), suggesting that V_D_ contributed much less to total genetic variance than V_A_.

We could directly estimate V_D_ for body size in the F2 cross and can thus provide data for this rarely estimated variance component. Although our V_D_ estimates for size were always associated with large SE, they suggest that V_D_ is present and environment dependent. The highest estimates of V_D_ were found in Dry and Hot for both male and female size, while it remained close to zero in Control and Hot-Dry. This environmental dependence of V_D_ has not been studied in detail before, although the environmental dependence of inbreeding depression was previously described (Bijlsma et al. 1999; Armbruster and Reed 2005; Fox and Reed 2011). Both V_D_ and inbreeding depression are expected to increase with inbreeding, for instance in shrinking populations when allele and genotype frequencies change because of increased drift (Falconer and Makay, 1996). This could have important implications since environmental changes may emphasize the effects of inbreeding on survival and thus on population size too, directly affecting genetic variances.

The large uncertainty around our estimates of V_D_ may come from the double-first cousin (DFC) breeding design used in the F2, despite the large sample size we had (827 family pairs with DFC offspring). Although the DFC design was proposed to estimate V_D_ (Fairbairn and Roff 2006), it may have limited statistical power because the probability that DFC share alleles identical by descent at a given locus is only 6.25 %, resulting in large SE, in contrast to full-sibs where this probability is 25%. Therefore, comparing maternal half-sibs and full-sibs might be a much more powerful approach to estimate V_D_, while it allows us to disentangle non-additive from maternal effects at the same time.

Although we could estimate genetic correlations between fitness and size with our data set (Supplemental Table S4), it was not possible to get unbiased estimates of those correlations. With our design, we could not rule out a strong confounding effect of population density because body size was measured in the F2, the offspring of beetles used for the fitness assay (Supplement Figure S1). Consequently, offspring of females with a high fitness (i.e. high offspring number) grew at a higher density since we used identical tubes with the same amount of flour for all females in the fitness assay.

#### Conclusions

Our study showed an increased adaptive potential in stressful conditions and a corresponding adaptation to those conditions after 20 generations. Although precise predictions of relative increase in fitness were not possible over this time period, we could make correct qualitative predictions. We expected and observed the highest relative fitness increase in the most stressful hot-dry condition and similar increases in single stress treatments. The apparently high adaptive potential of female beetles in Hot was limited by a negative genetic correlation with male fitness. We further found that genetic effects of males on fitness can be large and can increase the adaptive potential. Comparing genetic variances of fitness and size showed that they differed in their variance composition and in their cross-sex and cross-environment genetic covariances. Environmental effects on genetic variances were also not consistent between the two traits. We thus advise caution if studies interested in fitness V_A_ and the adaptive capacity of a population use body size as a proxy. Overall, we found that genetic variance in fitness is a key estimate of a population’s adaptive capacity for time scales over 20 generations. As such, it may help predict the adaptive response of populations exposed to new environmental conditions and help identify the populations most at risk of extinction. However, the reliability of such predictions will depend on the fitness estimate chosen and on the full integration of the multifaceted aspects of adaptation. Inclusion of genetic covariances between female and male fitness and genotype by environment interactions is thus important.

## Author contributions

FG and ELK designed the experiment. ELK and SHS conducted the experiment (crossing and fitness assay). SHS performed size measurements. ELK analysed the data. FG and ELK wrote the manuscript. SHS contributed and commented to manuscript.

## Acknowledgements

This work was supported by the Swiss National Science Foundation, grants PP00P3_1144846 and PP00P3_176965 to FG. We would like to thank all people helping with fitness measurements: Julian Bauer, Valérian Zeender, Maria Domenica Moccia, Sara Meier, Cilgia Lippuner, Tim Schoch, Tim Emmenegger, Elke Karaus.

