## Supplemental Information for "Genetic variance in fitness and its cross-sex covariance predict adaptation during experimental evolution"

**Contents**

**Figure S1:** Experimental design: Crossing design to produce paternal half-sib families and double-first cousins

**Figure S2**: Experimental Evolution design

**Figure S3:** Correlation abdominal segment IV Size with Prosternum Size

**Figure S4:** Distribution offspring number

**Figure S5**: Model diagnostic plots for analysis of genetic variance of offspring number

**Figure S6:** Model diagnostic plots for analysis of offspring number in control and selection lines after experimental evolution

**Figure S7:** Pairwise cross-environment genetic correlations of size

**Table S1:** Effect of condition on size. Results of linear mixed models

**Table S2:** Effect of selection regime on offspring number under different conditions after experimental evolution. Results of linear mixed models

**Table S3:** Genetic variances of size in female and male flour beetles and cross-sex additive genetic covariances and correlations in different environmental conditions when non-additive genetic effects were not included in the model

**Table S4:** Additive genetic correlations between fitness and size of females and males

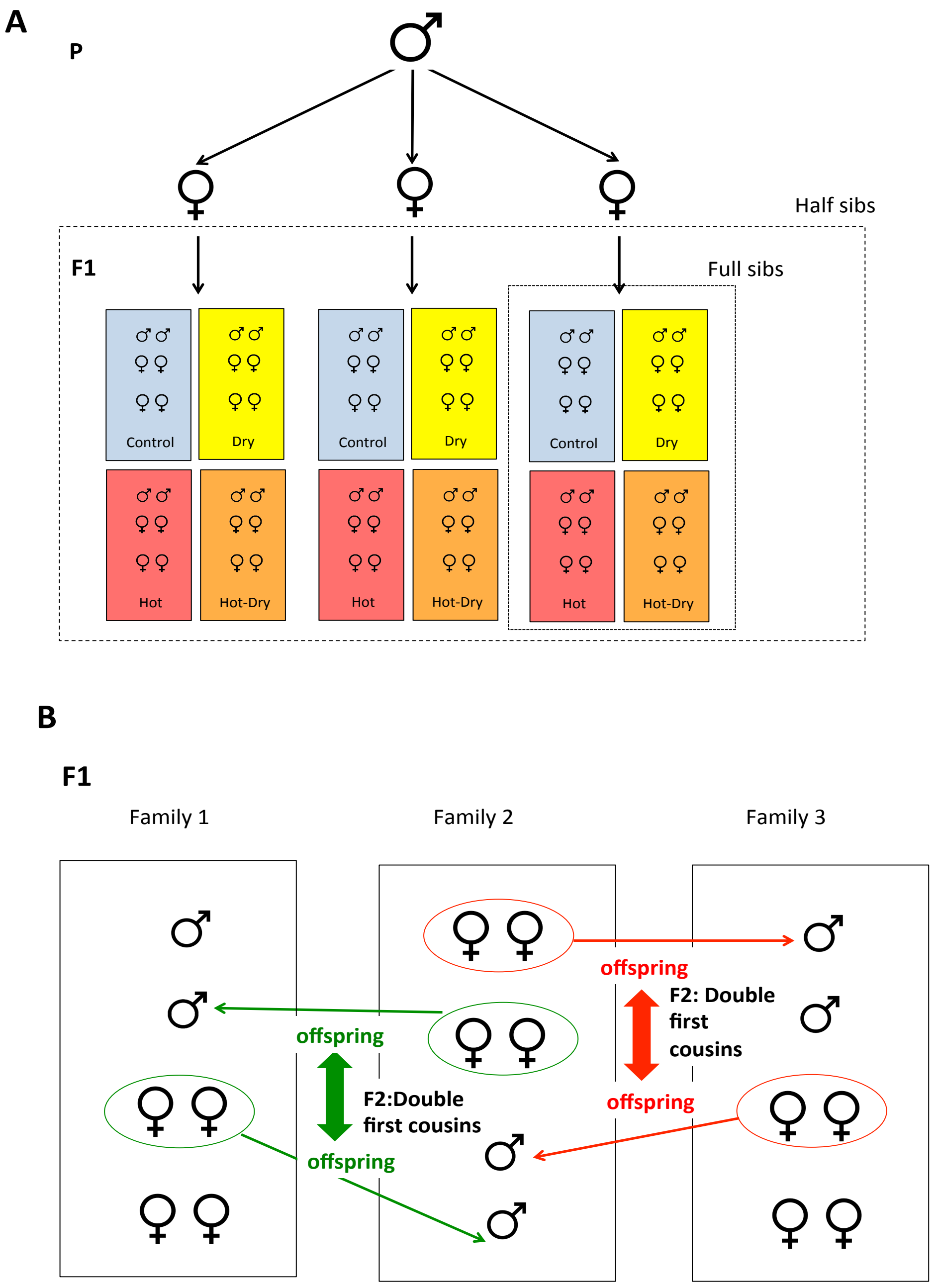


**Figure S1:** Crossing design. To produce paternal half-sib families, one male was mated subsequently with three females. Eggs of each female were randomly distributed across four different environmental conditions. Males and females of the next generation (F1) were used to estimate genetic variance in fitness **(A).** When we mated individuals of F1 to get offspring (used as our fitness estimate), we followed a specific crossing design. We always crossed two full-sib families (from different half-sib families) reciprocally. Offspring (F2) resulting from these two crosses are double first cousins **(B)**. F2 individuals were used to estimate additive and non-additive genetic variances in size.

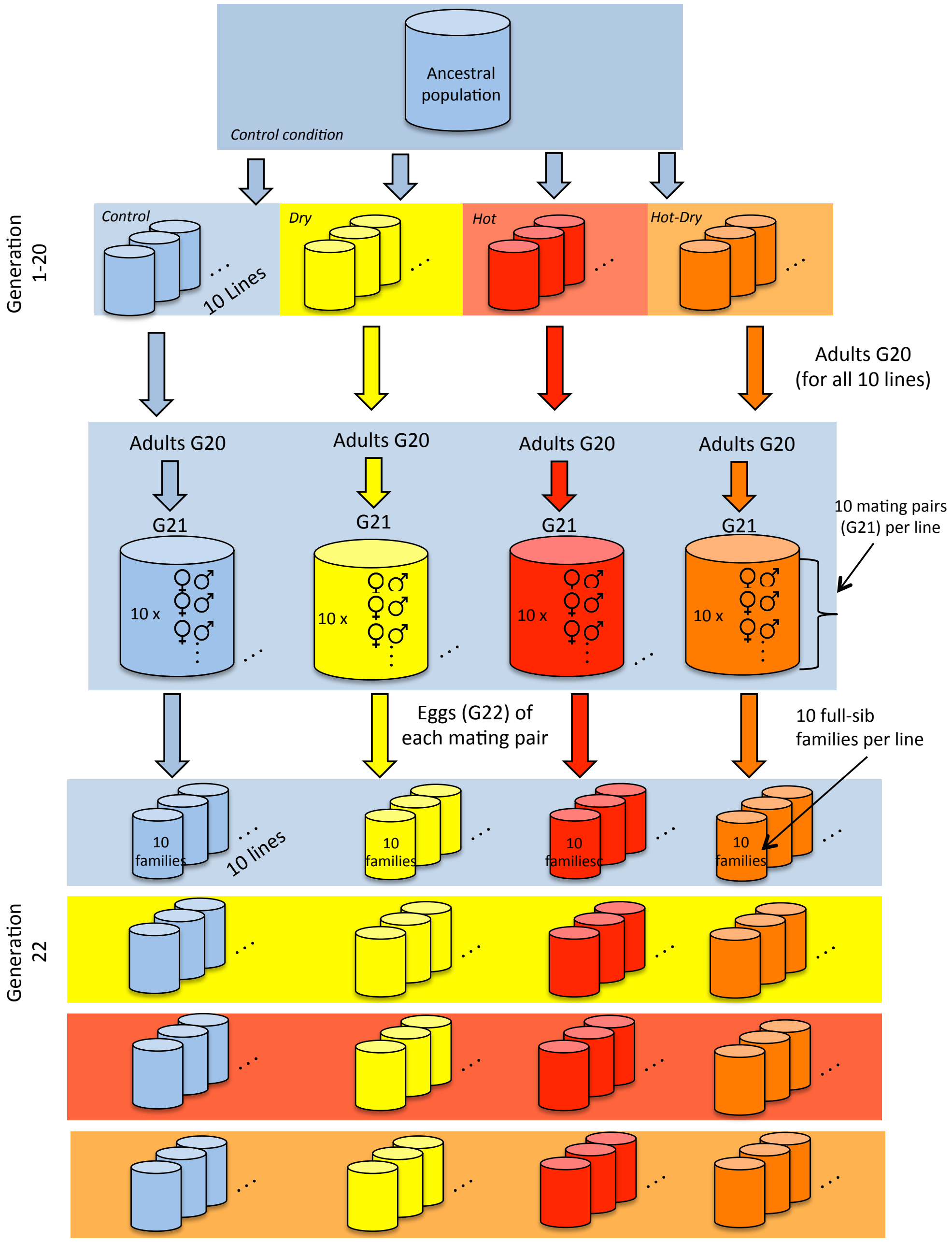


**Figure S2:** Schematic representation of the experimental evolution design. Lines were derived from a single outbred Tribolium castaneum population (collected in 2013, strain Cro1) that had lived under control conditions for ca. 30 generations. For each selection regime we used ten replicate lines. Lines stayed in control or treatment conditions for 20 generations. After one generation in control condition to remove potential maternal and epigenetic effects, individuals from all lines (generation 22) were transferred to control and treatment conditions in the egg stage. Fitness was measured in adult beetles by counting the number of adult offspring. Control conditions: 33°C, 70% relative humidity r.h.; Treatments: Dry: 33°C, 30% r.h.; Hot 37°C, 70% r.h.; Hot-Dry: 37°C, 30% r.h.

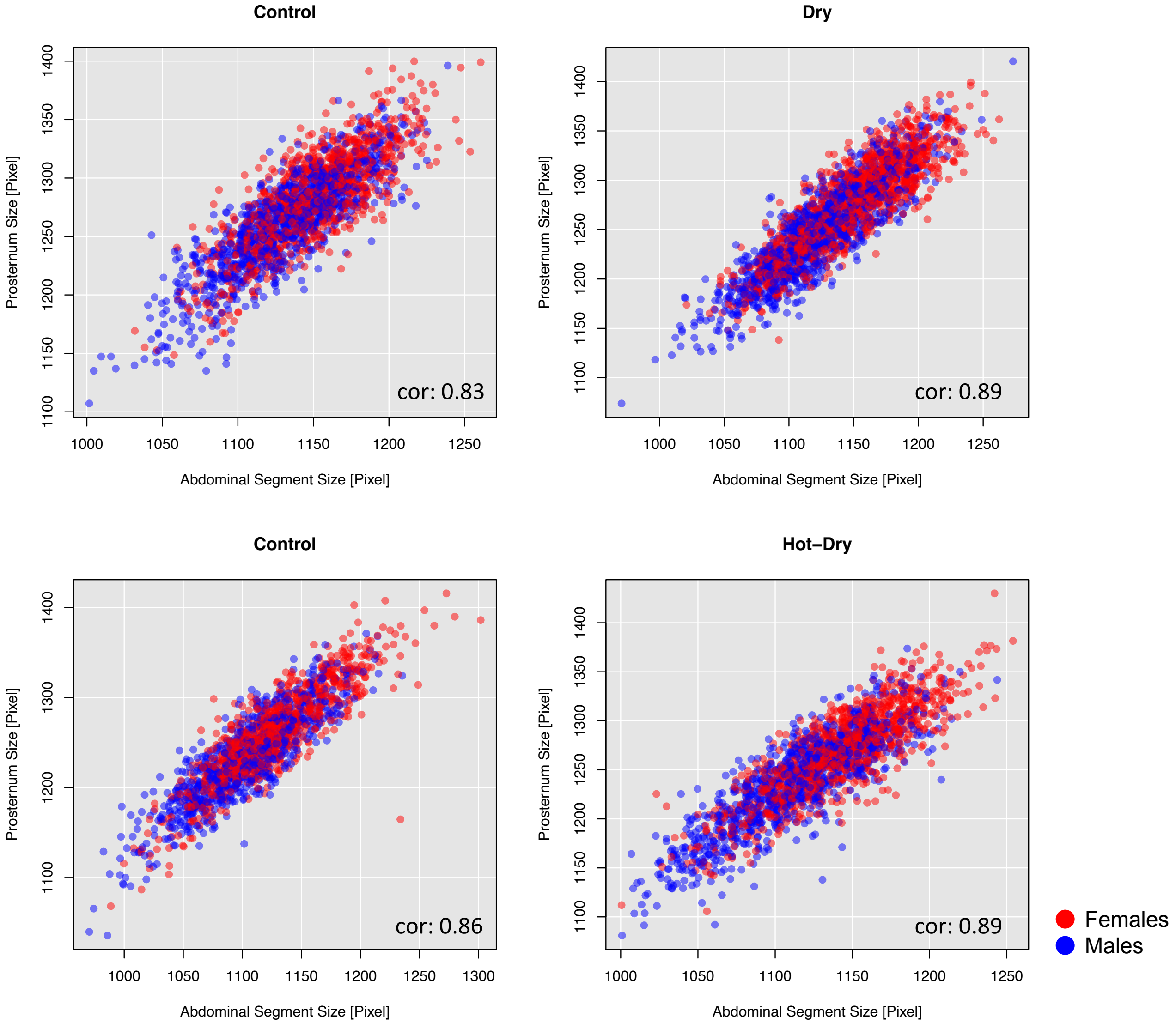


**Figure S3:** Correlation between Abdominal Segment Size IV and Prosternum Size in the four conditions

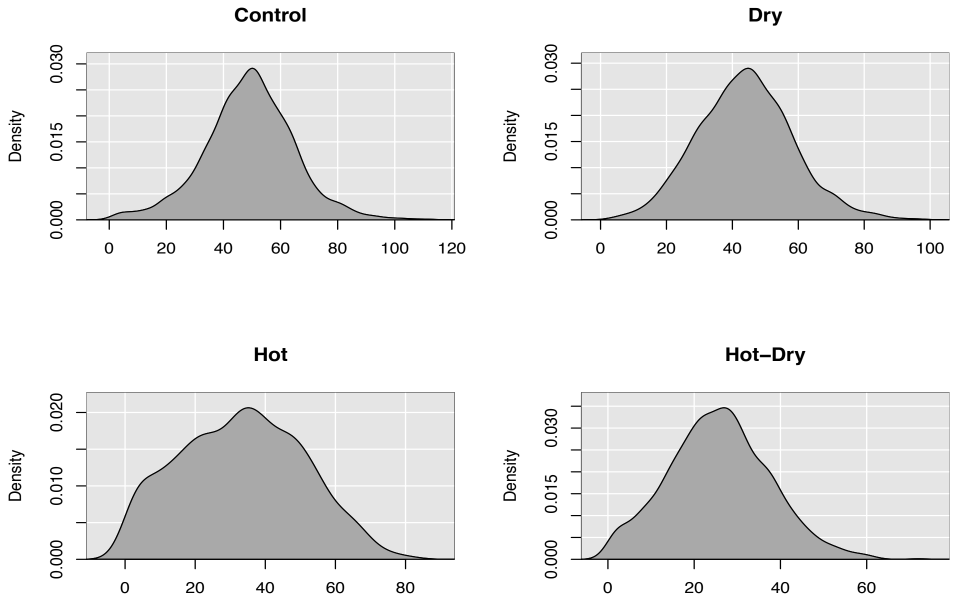


**Figure S4:** Distribution of offspring number in the four conditions

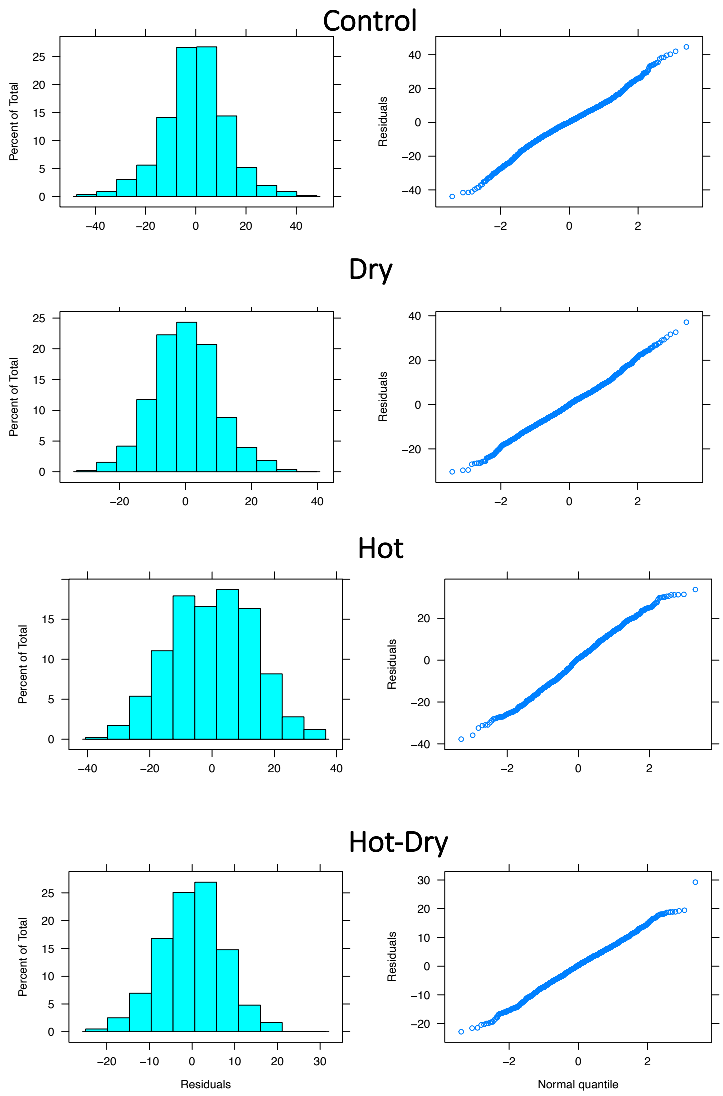


**Figure S5:** Model diagnostic plots (distribution residuals, qqPlot) for the analysis of genetic variance of fitness in four different conditions. A univariate animal model including additive genetic effects (breeding values) of females and males with a potential correlation between them was used to estimate additive genetic variance of fitness in females and males . $offspring_i = \mu + batch + a_{♀\,i} + a_{♂\,i} + e_i$ , where Offspring number of a female (i) depends on its additive genetic effects $a_{♀\,i}$ (its breeding value) and of additive genetic effects of the mating partner $a_{♂\,i}$ where $\mu$ is the population mean and $e$ a residual term. We included batch (day of starting the fitness assay) as fixed effect. Results are shown in Table 1 in the main manuscript.

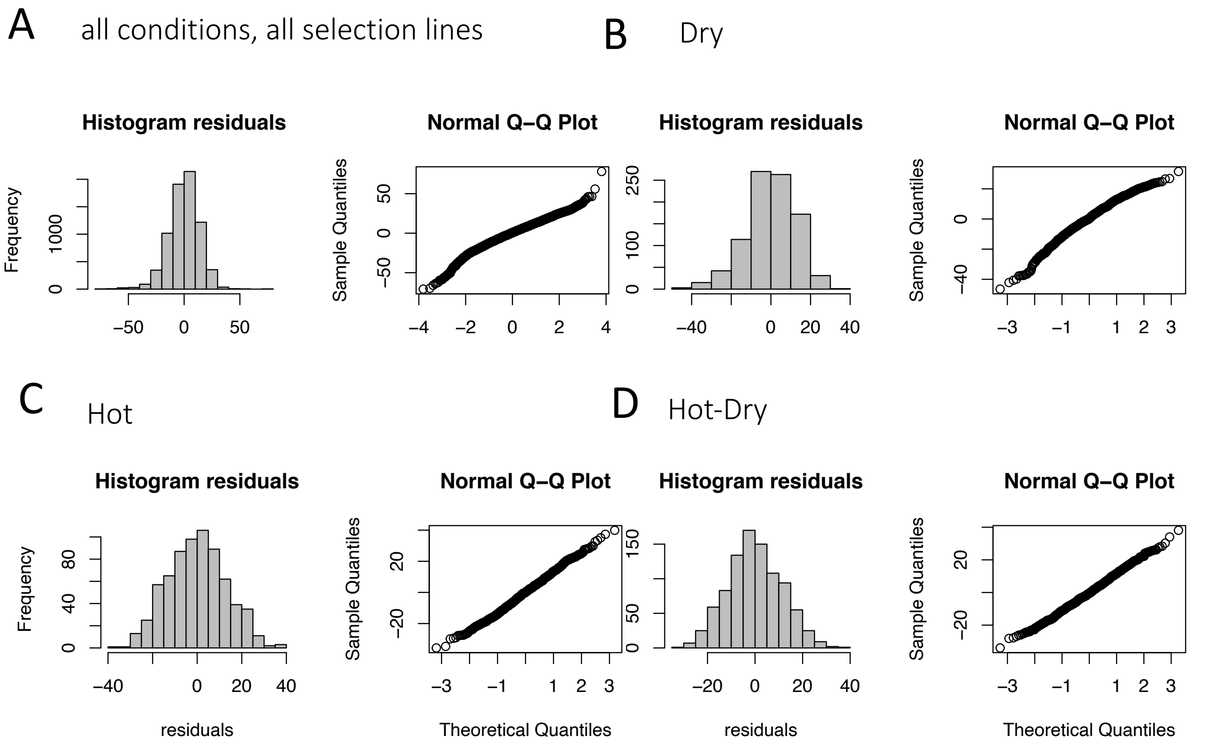


**Figure S6:** Model diagnostic plots for analyses of offspring number after experimental evolution. **A**: Model for testing whether adapatation to one condition increased fitness in other conditions. For this we run one analysis using the complete data set. Selection regime, conditions, and their interaction were used as fixed effects, lines, families (implicitly nested within lines), and line-treatment interaction as random effects : Offspring ~μ+Selection*Treatment + *line*+ *line**Treatment+ *family* +*e,* where μ is the mean, Selection the selection regime, Treatment the condition in which fitness (=offspring number) was measured, selection line and family of each individual are included as random effects and *e* is the error deviation. **B-D**: Separate models to test adaptation to each condition using only data from control lines and native (e.g. dry lines in dry ) selection lines with Selection as fixed effects and lines and family as random effects. **B** dry conditions; **C**: hot conditions; **D**: hot-dry conditions.

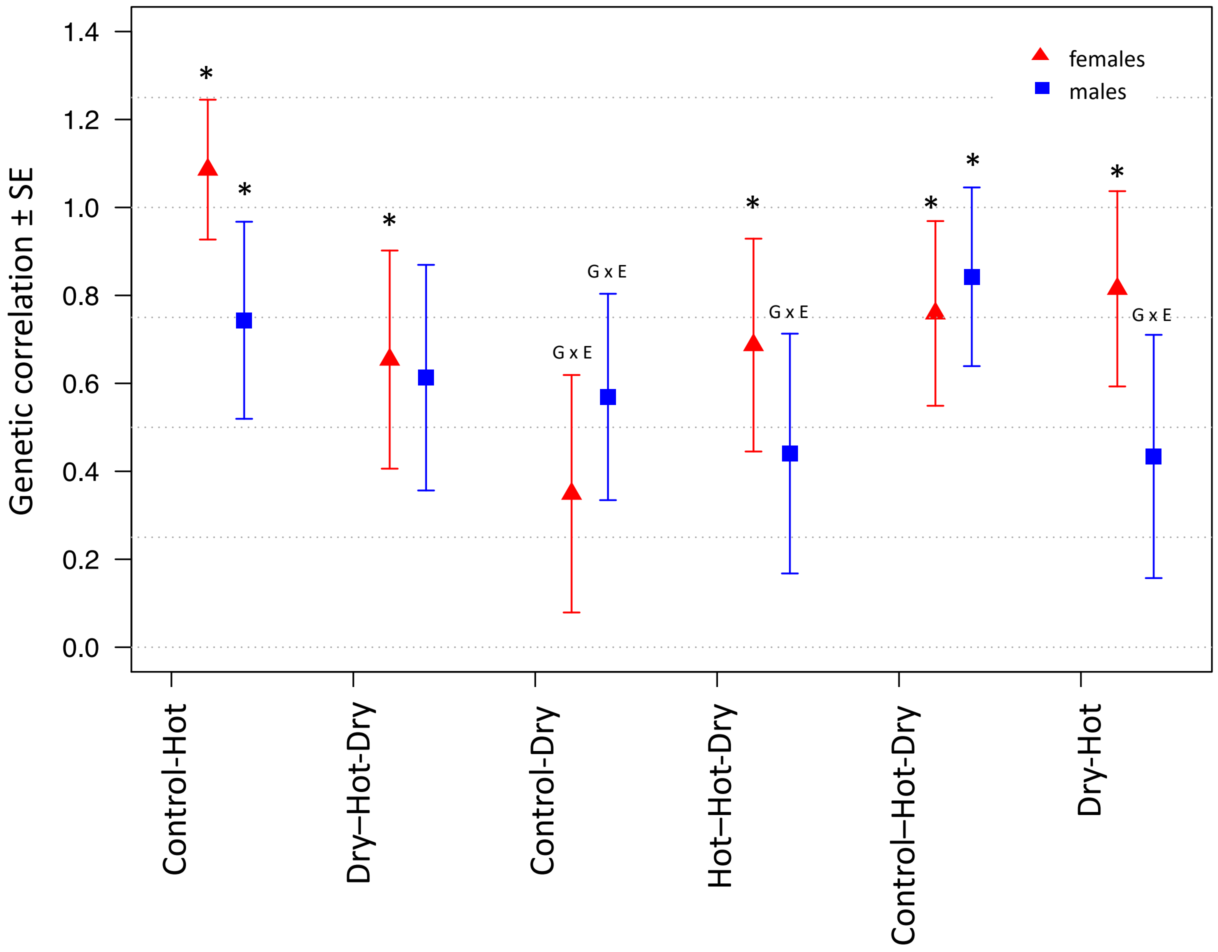


**Figure S7:** Pairwise cross-environment genetic correlations of size (centroid size of abdominal segment IV) in *Tribolium castaneum* estimated using bivariate animal models. * show significant correlations, i.e. significantly different from zero. G x E (= genotype by environment interactions) indicate correlations significantly different from one. Control: 33°C, 70% relative humidity (r.h.); Dry: 33°C, 30% r.h.; Hot: 37°C, 70% r.h.; Hot-Dry: 37°C, 30% r.h. To achieve model convergence for estimating genetic correlations between female size in Control and Hot we had to use a model with a completely unstructured and unconstrained genetic covariance matrix, which resulted in a correlation >1.
